# Deficits in neural encoding of speech in preterm infants

**DOI:** 10.1101/2023.05.09.539966

**Authors:** Nikolay Novitskiy, Peggy H. Y. Chan, Mavis Chan, Chin Man Lai, Tak Yeung Leung, Ting Fan Leung, Marc H. Bornstein, Hugh S. Lam, Patrick C. M. Wong

## Abstract

Preterm children show developmental cognitive and language deficits that can be subtle and sometimes undetectable until later in life. Studies of brain development in children who are born preterm have largely focused on vascular and gross anatomical characteristics rather than pathophysiological processes that may contribute to these developmental deficits. Neural encoding of speech as reflected in EEG recordings is predictive of future language development and could provide insights into those pathophysiological processes. We recorded EEG from 45 preterm (≤ 34 weeks of gestation) and 45 full-term (≥ 38 weeks) Chinese-learning infants 0 to 12 months of (corrected) age during natural sleep. Each child listened to three speech stimuli that differed in lexically meaningful pitch (2 native and 1 non-native speech categories). EEG measures associated with synchronization and gross power of the frequency following response (FFR) were examined. ANCOVAs revealed no main effect of stimulus nativeness but main effects of age, consistent with earlier studies. A main effect of prematurity also emerged, with synchronization measures showing stronger group differences than power. By detailing differences in FFR measures related to synchronization and power, this study brings us closer to identifying the pathophysiological pathway to often subtle language problems experienced by preterm children.

## 1. Introduction

The present study investigates neural encoding of speech in preterm and full-term infants, with a goal to understand the neural functional underpinnings of preterm language and communication deficits. Such an understanding holds the potential for developing objective, early detection tools for language impairments that affect as much as one-third of the preterm population and are of continuing concern in cognitive development and education(Putnick, Bornstein, Eryigit-Madzwamuse, & Wolke, 2017; Sansavini et al., 2010; Woodward et al., 2009).

Preterm birth is an important clinical condition with a prevalence of 11% of all births (Blencowe et al., 2012; Walani, 2020). By school age, preterm-born children score consistently lower than full-term children in many areas of language and literacy (Luu et al., 2009; Zimmerman, 2018), including complex language features such as formulating sentences (van Noort-van der Spek, Franken, & Weisglas-Kuperus, 2012). Importantly, even children born moderate-to-late preterm have poorer long-term neurodevelopmental outcomes (McGowan, Alderdice, Holmes, & Johnston, 2011). Children who are born late preterm are at greater risk for language difficulties than for other neurodevelopmental disorders such as ADHD (Harris et al., 2013; Rabie, Bird, Magann, Hall, & McKelvey, 2015). These deficits in language and communication are likely associated with differences in the central nervous system. Brain studies of preterm infants historically focused on gross pathologies, such as periventricular leukomalacia and intraventricular haemorrhage (Guit, van de Bor, den Ouden, & Wondergem, 1990; Kidokoro et al., 2014; van de Bor, den Ouden, & Guit, 1992), whereas recent studies have used MRI to investigate more detailed anatomical measures, such as fractional anisotropy (FA), to differentiate full-term and preterm groups (Brignoni-Pérez et al., 2022). FA increases linearly during language learning over 9 months (Schlegel, Rudelson, & Tse, 2012), and animal data have revealed a correlation between FA changes and myelin increase in histological follow-ups (Almeida & Lyons, 2017). Preterm infants are more likely to have smaller brain size (Bouyssi-Kobar et al., 2016; Walsh, Doyle, Anderson, Lee, & Cheong, 2014), reduced gray matter volume (Munakata et al., 2013), abnormal white matter microstructure (Dibble, Ang, Mariga, Molloy, & Bokde, 2021; Kelly et al., 2019; Thompson et al., 2019), delayed gyral maturation (Walsh et al., 2014), and lower neurite density (W. Wang et al., 2022). These structural measures, however, do not address differences in brain function, nor can behavioural patterns be attributed to physiological processes. Even with functional measures, MRI cannot capture millisecond-level changes in the speech signal that define spoken language.

Speech encoding is foundational to language development. Minute millisecond-by-millisecond changes in speech can result in changes in word meaning (for example, altering the pitch patterns of /ma/ can result in four different words in Putonghua Chinese). Infants as young as 4 months can differentiate minimal differences in the speech signal (Eimas, Siqueland, Jusczyk, & Vigorito, 1971; Polka & Werker, 1994). EEG, with its sub-millisecond resolution, provides the needed temporal precision to examine neural encoding of speech. Specifically, the EEG-derived Frequency Following Response (FFR) is a potential biomarker of a range of developmental language and communication problems (Font-Alaminos et al., 2020; Ribas[Prats et al., 2022). FFR belongs to a family of mostly (but not exclusively) brainstem-originating potentials of which the best known representative is the click-evoked auditory brainstem response (ABR; Skoe & Kraus, 2010). ABR peaks demonstrate delayed latencies in preterm as compared to full-term infants (Ribeiro & Carvallo, 2008; Silva, Lopez, & Mantovani, 2014; Sleifer et al., 2007). A meta-analysis of 14 studies demonstrated a large and significant effect size for the latency difference between preterm and full-term groups in a III – V ABR peak latency difference (Stipdonk et al., 2016). However, the click response still does not provide the kind of precision needed to forecast the development of linguistic skills in individual children compared to actual speech (Wong et al., 2021). Earlier studies demonstrated a dissociation between ABR and FFR parameters in individual adults (Hoormann, Falkenstein, Hohnsbein, & Blanke, 1992). Furthermore, the click response is not sufficiently rich in the spectral domain to operationalise underlying pathophysiology. Thus, contemporary studies have used natural, spectrally rich speech sounds to elicit spectrally rich FFR that could be decomposed into a larger set of physiologically meaningful measures. For example, Novitskiy *et al*. (2022) reported a rapid progression of different measures of FFR to speech during the first 2 years of life, and Wong *et al*. (2021) found that FFR to speech sounds during infancy predicted language learning in toddlerhood.

The scalp-recorded FFR to speech represents a direct neural mirroring of perceived harmonic sounds in speech signals (Anderson, Parbery-Clark, White-Schwoch, & Kraus, 2015; Arenillas-Alcón, Costa-Faidella, Ribas-Prats, Gómez-Roig, & Escera, 2021; Jeng, Lin, & Wang, 2016; Kraus & White-Schwoch, 2020; Krizman & Kraus, 2019). Generally speaking, FFR measures can be classified into those related to synchronization (phase-locking) to the stimulus and those related to signal power. For example, inter-trial phase coherence (ITPC) directly measures phase-locking in EEG (Tallon-Baudry, Bertrand, Delpuech, & Pernier, 1996). By contrast, the residual root-mean-squared noise (Noise RMS) in the pre-stimulus interval after averaging EEG trials indicates the power of spontaneous activity (Skoe, Krizman, Anderson, & Kraus, 2015). In the present study, we compared FFR to speech in preterm versus full-term infants. Our goals were to investigate whether preterm infants show poorer neural encoding of speech relative to their full-term peers and to determine whether FFR measures that primarily indicate deficiencies in either synchronization (e.g., Pitch Strength) or power (e.g., Noise RMS) are differentially affected by preterm birth. This study represents one of the first attempts to identify specific pathophysiological processes underlying the functional language deficit associated with preterm birth.

We expected that different FFR measures (dependent variables) would relate to synchronization versus gross power on a Specificity Principle. This principle postulates that specific processes in specific individuals are affected by specific experiences at specific time periods (Bornstein, 2019; Lerner & Bornstein, 2021). In our case, the setting of each specific infant is a specific transition from *in utero* to *ex utero* environments at a specific time after gestation, which affects two specific processes, synchronization and power growth in a specific way as reflected by our dependable variables. We expected that this coaction would result in specific changes in speech development later in life, as the same dependable variables have been shown to predict speech after 1 year of postnatal development (Wong et al., 2021). Including multiple dependent variables in the study allowed us to detect specific and differentiating independent-dependent variable relations.

## 2. Materials and methods

### 2.1. Participants

Forty-five preterm infants (25 females) and 45 full-term infants (21 females) from homes where Cantonese is spoken as the dominant language participated. All infants were of Chinese ancestry. Preterm infants were born before 34 weeks of gestation, and full-term infants were born after 38 weeks of gestation (Committee on fetus and newborn, 2004; The American College of Obstetricians and Gynecologists Committee, 2013). Median gestational age at birth was 31(7) weeks for preterm infants and 38(1) weeks for the full-term infants (parentheses enclose interquartile ranges, i.e. the difference between the 25th and 75th percentiles of the distribution). Median birth weights were 1.48(1.01) kg and 3.06(0.41) kg in preterm and full-term groups, respectively. The corrected age of a preterm infant was calculated by subtracting the number of weeks between the birth and expected date of delivery (40 weeks) from their chronological age (Committee on fetus and newborn, 2004). At the time of EEG data acquisition, the median corrected age of the preterm infants was 4(4) months, and the median chronological age of the full-term infants was also 4(3) months (Figure 1A). Median maternal education was 16(5) years and 16(4) years in preterm and full-term groups (Figure 1B). Maternal education was used as a proxy for SES, following previous research (e.g., Bornstein, Hahn, Suwalsky, & Haynes, 2003; Poulsen et al., 2019; see McHale et al., 2022, for a review). The success rates of EEG testing were 95% and 86% respectively, for preterm and full-term infants.

**Figure 1.**
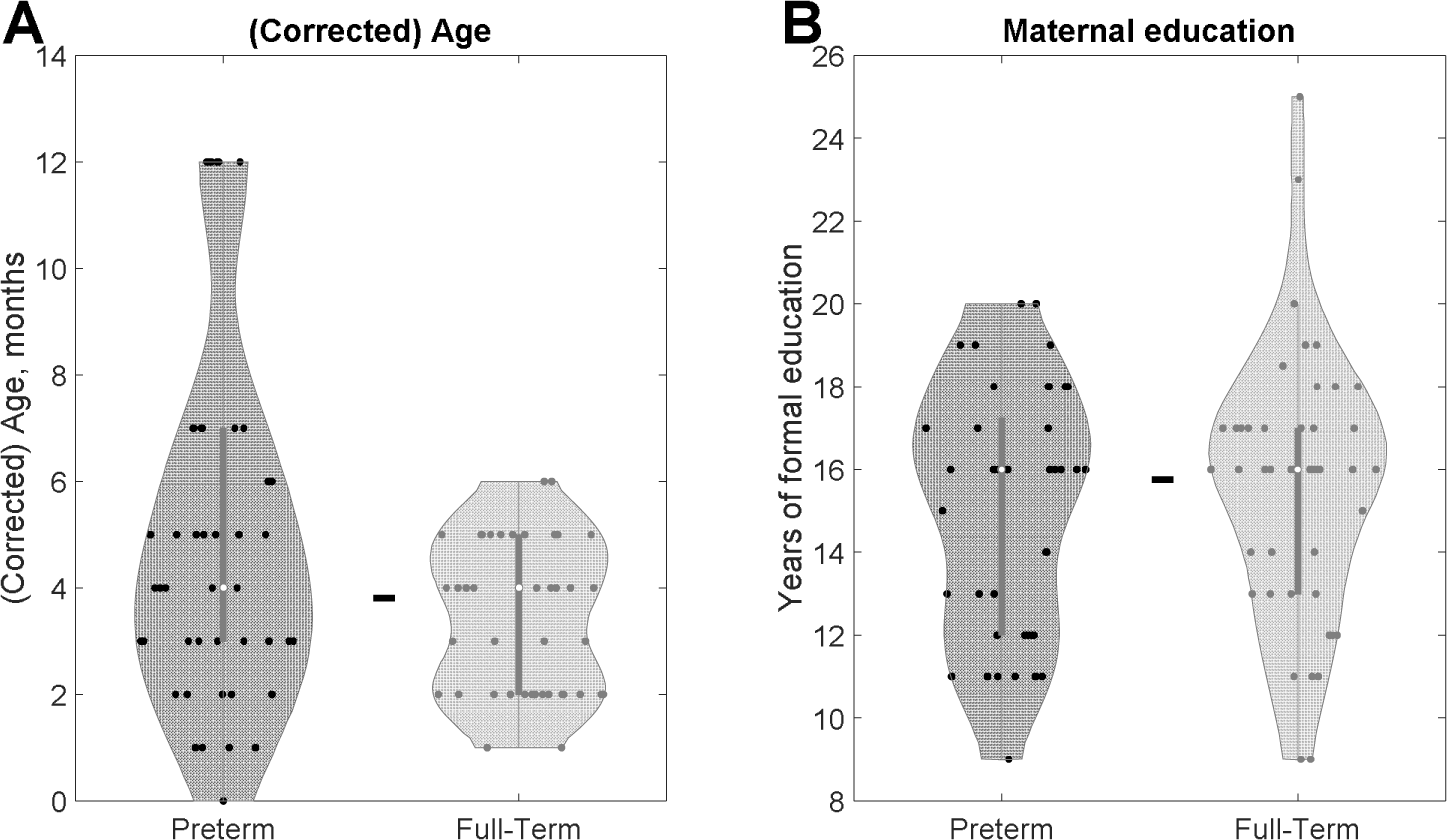
The distributions (corrected) age (A) and maternal education (B) in the preterm and full-term groups. No significant group difference was detected in either measure, as indicated by the dash between the violins.

### 2.2. Procedures

Infants were held by their principal caregiver and were naturally asleep during the recording. Recording was conducted inside a soundproof and electrically shielded room. A random sequence of three equiprobable lexical tones embedded in the syllable /ga/ was presented to infants, 620 times each. The tones were female-voice Cantonese tone 2 (rising high) and tone 4 (falling low) and Putonghua tone 3 (dipping tone) that have been used in previous studies (Novitskiy et al., 2022; Wong et al., 2021). Sounds were presented by Neurostim (Compumedics) or E-prime-3 (Psychology Software Tools). The inter-stimulus interval (ISI) was 331 ms in 45 infants (21 preterms) and 501 ms in the remaining 45 infants (24 preterms), which resulted in stimulus-onset-asynchronies (SOA) of either 507 or 682 ms (χ*^2^* = 0.4, *P* = 0.53 for ISI type per experimental group).

### 2.3. EEG recording and analysis

Continuous EEG was recorded with the Neuroscan system (Compumedics, Australia) at 20 KHz sampling rate from Cz and left and right mastoids with Cpz as reference and Fpz as ground. The impedance was kept below 2 kOm. Offline analysis was performed on Cz which was re-referenced to the average of the two mastoids. Continuous EEG data were filtered with a zero-phase digital bandpass FIR filter (bandpass 80-1500 Hz, filter order 512, MATLAB functions *filtfilt* and *designfilt*). The 80-Hz low cutoff of the filter left out the non-target low-frequency activity such as long-latency responses (LLRs) to auditory stimuli, alpha-rhythm, and eye-blink artifacts, and the 1500-Hz high cutoff of the filter removed frequencies that fell above the phase-locking capacities of the FFR (Skoe & Kraus, 2010). Filtered data were epoched from -50 to 250 ms from stimulus onset with epochs exceeding ±25 μV rejected during the analysis. Averaged across epochs, FFR was analysed in both time and frequency domains and with time-frequency (TF) decomposition. The FFR power spectrum was calculated as the squared Fast Fourier transform (FFT) of the post-stimulus part of the averaged FFR. The resulting power was transformed into decibels (dB) relative to one millivolt (dBmV). The FFR ITPC (Inter-trial Phase Coherence) was calculated as an absolute value of the vector average across normalised FFTs of the single-epoch FFR. The ITPC TF decomposition (periodogram) was obtained by calculating ITPC in sliding 50-ms Hamming windows in 2-ms steps along the FFR epoch.

Altogether, 11 FFR parameters were extracted from the FFR (**Figure 2**):

1. Response Consistency was calculated as a Fisher-transformed averaged correlation between the FFRs from two halves of the randomly split single-epoch data after 300 iterations.
2. Low-Frequency Spectral Power was calculated as a mean over the 120-260-Hz interval of the FFR power spectrum. This range corresponds to the fundamental frequency (F0); that is, the first harmonic of female voice.
3. Mid-Frequency Spectral Power was calculated as mean over the 260-750-Hz interval of the FFR power spectrum. This range corresponds to the second and third harmonics of female voice.
4. Low/High Power Ratio was calculated as a difference between Low-Frequency Spectral Power (2) and Higher-Frequency Spectral Power. The latter was calculated as a mean over the 750-1200-Hz interval of the FFR power spectrum. Decibel (dB) values were log transformed where the log ratio between two values equals the difference between those values: log(a/b)=log(a)-log(b). Thus, for dB-transformed values a difference is equivalent of the ratio of the non-transformed values.
5. Mid/High Power Ratio was calculated as a difference between Mid-Frequency Spectral Power (3) and Higher-Frequency Spectral Power.
6. Noise root-mean square (RMS) was the RMS of the 50-ms pre-stimulus interval of the FFR waveform.
7. Mid-Frequency ITPC was calculated as the mean over 260-750-Hz interval of the whole-epoch FFR ITPC.
8. Low-Frequency ITPC was calculated as the mean over 120-260-Hz interval of the whole-epoch FFR ITPC.
9. Maximal ITPC was the highest value across the ITPC periodogram.
10. Signal-to-noise ratio (SNR) was calculated as the dB-transformed ratio of the root-mean-square (RMS) power of the 170-ms post-stimulus to 50-ms pre-stimulus interval of the FFR waveform.
11. Pitch Strength was calculated as the maximum of the FFR autocorrelation waveform excluding the initial peak at 0 ms delay.

**Figure 2.**
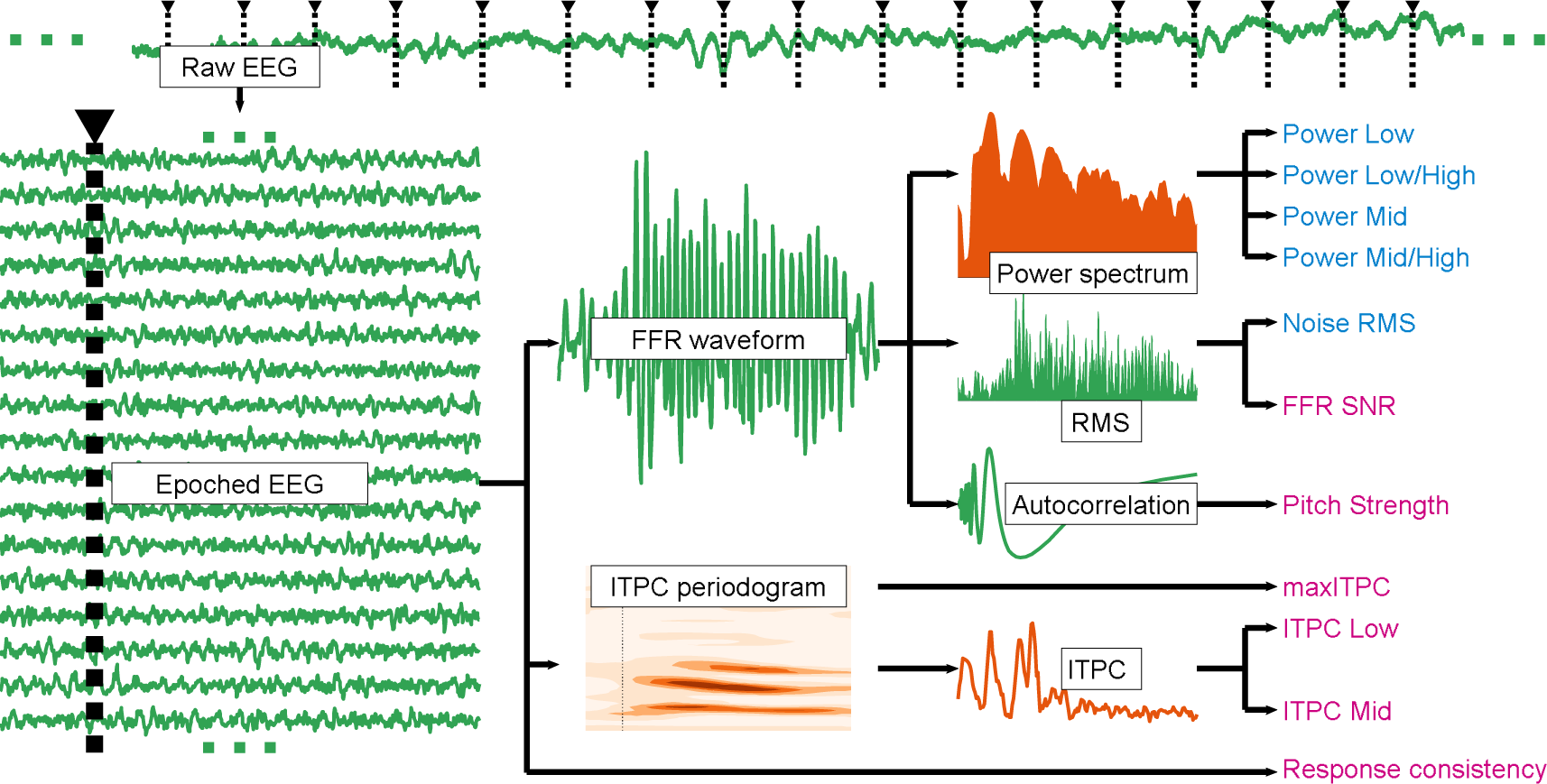
Two types of FFR measures: Synchronization and Power. Raw EEG signals (the upper panel) are recorded from the listener’s central nervous system, re-referenced, filtered, and aligned with the onsets of a particular external auditory stimulus (indicated by the inverted triangles with dashed lines) to produce the epoched EEG (left lower panel). EEG trials (epochs) are shown as rows at the epoched EEG panel. The ellipses frame Raw and Epoched EEG waveforms to indicate that they show a subset of a larger data set. Epoched EEG was further transformed into time-domain (green) and frequency-domain (orange) representations that included averaged FFR waveforms and ITPC periodograms, FFR Power Spectrum, Root-mean square (RMS) and Autocorrelation. These representations can be mapped onto Synchronization (purple) and Power (blue) FFR measures. See Discussion for a detailed explanation of the mapping. As stated in Results, our a priori classification converged with the statistical clustering results (see Figure 4).

These dependent variables represent a subset of parameters which were measured in a group of term infants without neurological abnormalities (Novitskiy et al., 2022) and included in a machine-learning model that successfully predicted speech outcomes (Wong et al., 2021). In the current study, we selected those dependent variables that can be measured fully objectively and whose estimation does not require knowledge about the auditory stimulus. For example, Pitch Error and Pitch Tracking are important measures of the brain’s ability to follow the pitch of the stimulus (Jeng, Peris, Hu, & Lin, 2013). However, calculations of these dependent variables require the extraction of pitch of the presented auditory stimulus which reduces their value as universal measures. Traditional peak latency and amplitude measurement provide a great deal of physiological information (Anderson, Parbery-Clark, White-Schwoch, & Kraus, 2012; Skoe et al., 2015) but require meticulous peak-picking by trained personnel which reduces their value potential for use in a mass-screening instrument.

### 2.4. EEG measure clustering

FFR measures can be classified into one of two types which reflect, respectively, either synchronization or gross power (Bourgeois, Jastreboff, & Rakic, 1989; Liu, Palmer, & Wallace, 2006; Seidl & Rubel, 2016). Specifically, ITPC, Pitch Strength, FFR SNR and Response Consistency can be considered synchronization-related measures because they reflect a degree of phase-locking between the stimulus and neural response in the brainstem (Jeng, Lin, Sabol, et al., 2016; Jeng, Lin, Chou, et al., 2016). Noise RMS, Spectral Power, and Power Ratio can be regarded as measures of gross power because they reflect background EEG noise and activity (Skoe et al., 2015). To obtain converging evidence for this process-de d grouping, we employed a data-driven approach that assessed statistical relations and clustering patterns of the FFR measures extracted in this study.

To do so, we first performed principal component analysis (PCA) of the 11 FFR measures. The resulting data matrix was composed of 270 rows (90 participants x 3 tones) and 11 columns (measures). The data were normalised across participants and tones with the MATLAB *zscore* function. PCA was performed with the MATLAB *pca* function. The outcome was 11 principal components with their coefficients (loadings), scores, and the percentages of variance explained. We selected the first two components on the Kaiser rule (i.e., the components with eigenvalues higher than unity). They explained 79.6% of the variance with 43.9% explained by the first component (PC1) and 35.7% explained by the second component (PC2). Loadings of the PC1 and PC2 were rotated with varimax rotation to align the maximal variance with the component axes.

We then performed k-means clustering in two clusters of the first two principal component coefficients of the measures. We used the MATLAB function *k-means* with default parameters (the squared Euclidean distance metric and the k-means++ algorithm for cluster center initialization) which was replicated five times to achieve a stable solution (Cohen, 2017). The number of clusters was set to two according to our *a priori* hypothesis that postulated two essential neuronal characteristics, synchronization and gross power. The clustering of the FFR measures in the principal component space converged with our *a priori* bipartite classification being primarily related to neural synchronization or gross power (henceforth Synchronization and Power variables, respectively). The scores of the two principal components were entered into the statistical analysis. In addition, we performed statistics on the individual measure values. See Results below for details.

### 2.5. Statistical analysis

The scores of the two principal components and individual measures were fed into a 4-way fixed-effects ANCOVA (MATLAB function *fitlm* with method *anova*) with two discrete factors of Prematurity (preterm vs. full-term) and Tone (ga2, ga3, or ga4) and two continuous factors of Corrected Age at FFR data collection (in months) and Maternal Education (in years). False-discovery rate (FDR) correction for multiple comparisons was applied whenever necessary (Benjamini & Hochberg, 1995). Age at FFR data collection and Tone need to be accounted for because significant effects of these measures have been found in previous studies (Novitskiy et al., 2022; Wong et al., 2021). Gestational age and birth weight have also emerged as significant factors in previous studies. However, because Prematurity is already accounting for gestational age and correlating with birth weight, these two factors were not additionally examined in the present study. Maternal Education is a component of socioeconomic status (SES) that was previously shown to affect the risk of premature birth in European cohorts (Ruiz et al., 2015). We included it in the model even though SES did not improve language outcome to prediction in a previous study of Chinese children in Hong Kong (Wong et al., 2021).

### 2.6. Data availability

The data that support the findings of this study are available from the corresponding author upon reasonable request.

## 3. Results

A Chi-squared test revealed no reliable difference between groups in sex ratios (χ*^2^ =* 0.71*, P =* 0.399) or EEG success rates (χ*^2^* = 0.167, *P* = 0.68). Birth weight correlated with gestational age (Spearman ρ *=* 0.845*, P =* 1.23·10^-25^); expectedly, preterm infants weighed less at birth (Wilcoxon test*, Z =* -7.71*, P =* 1.3·10^-14^). A Wilcoxon test revealed no between-group difference in chronological/corrected age (*Z =* 1.85*, P =* 0.064, Figure 1A). Median maternal education did not differ between the two groups (Wilcoxon test, *Z* = -0.49, *P*=0.62, **Figure 1B**).

Preterm and full-term infant grand-averaged FFR waveforms for the three tones are shown in **Figure 3**. Visual inspection confirms that the FFR waveforms of the preterm infants are noisier than those of their full-term peers.

**Figure 3.**
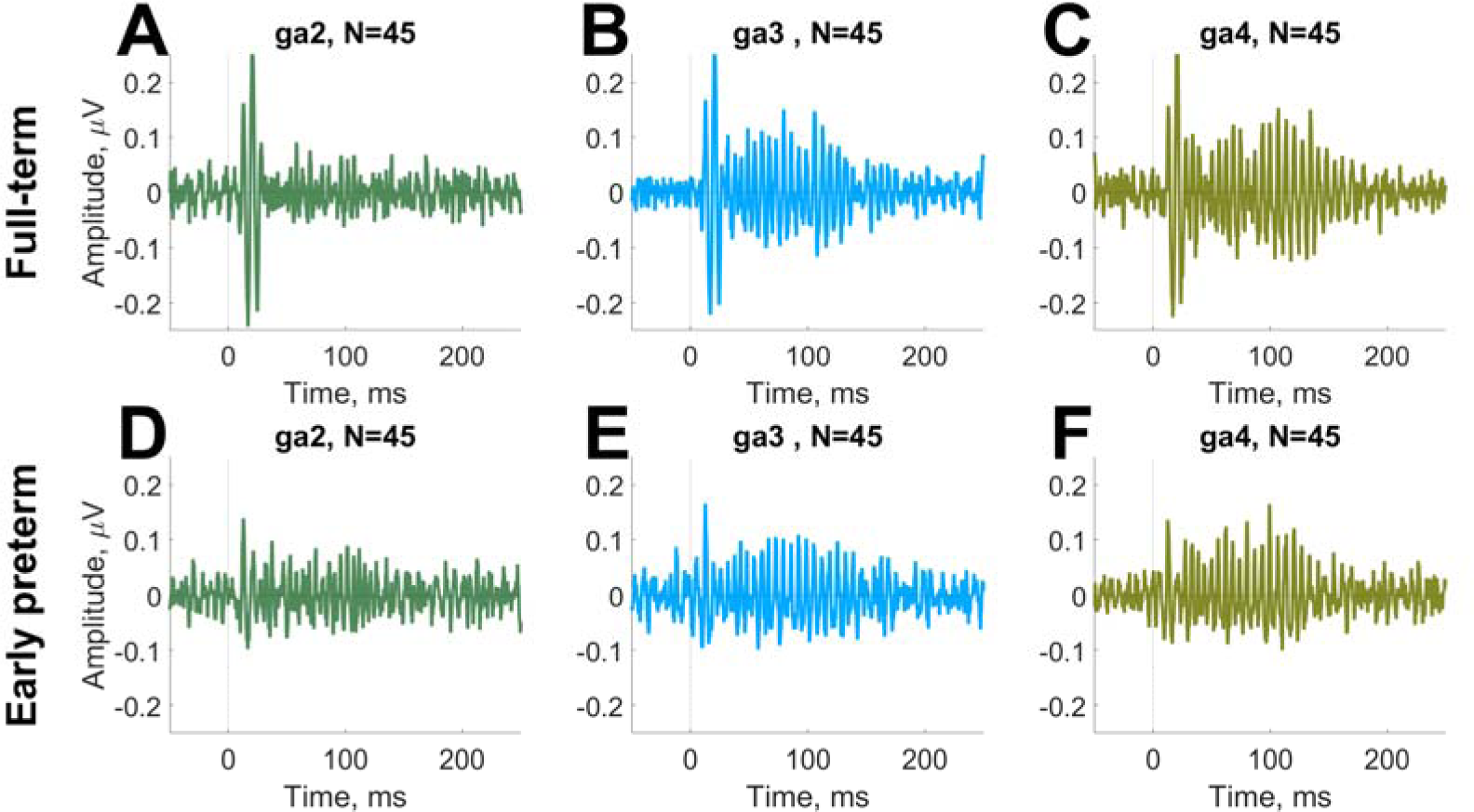
The grand-averaged FFR responses of the preterm (**D-F**) and full-term (**A-C**) infants to the three speech stimuli: /ga2/ (**A, D**), /ga3/ (**B, E**), and /ga4/ (**C, F**).

**Figure 4A** illustrates the results of the k-means clustering. The PC coefficients (loadings) for each measure are plotted as open circles connected with the lines to the centroid of their cluster that is plotted as a filled circle. The best total sum of squared Euclidian distances (i.e., minimal of the five replications) was 0.0222, and it replicated five out of five times. Coordinates of the centroids were [0.4056, 0.0002] for Synchronization and [-0.0001, 0.4452] for Power, where first coordinate is the PC1 value and second coordinate is the PC2 value. The sum of squared Euclidian distances between the measures of a cluster and cluster centroid was 0.0117 for Synchronization and 0.0105 for Power, while the distance between the centroids was 0.3626.

**Figure 4.**
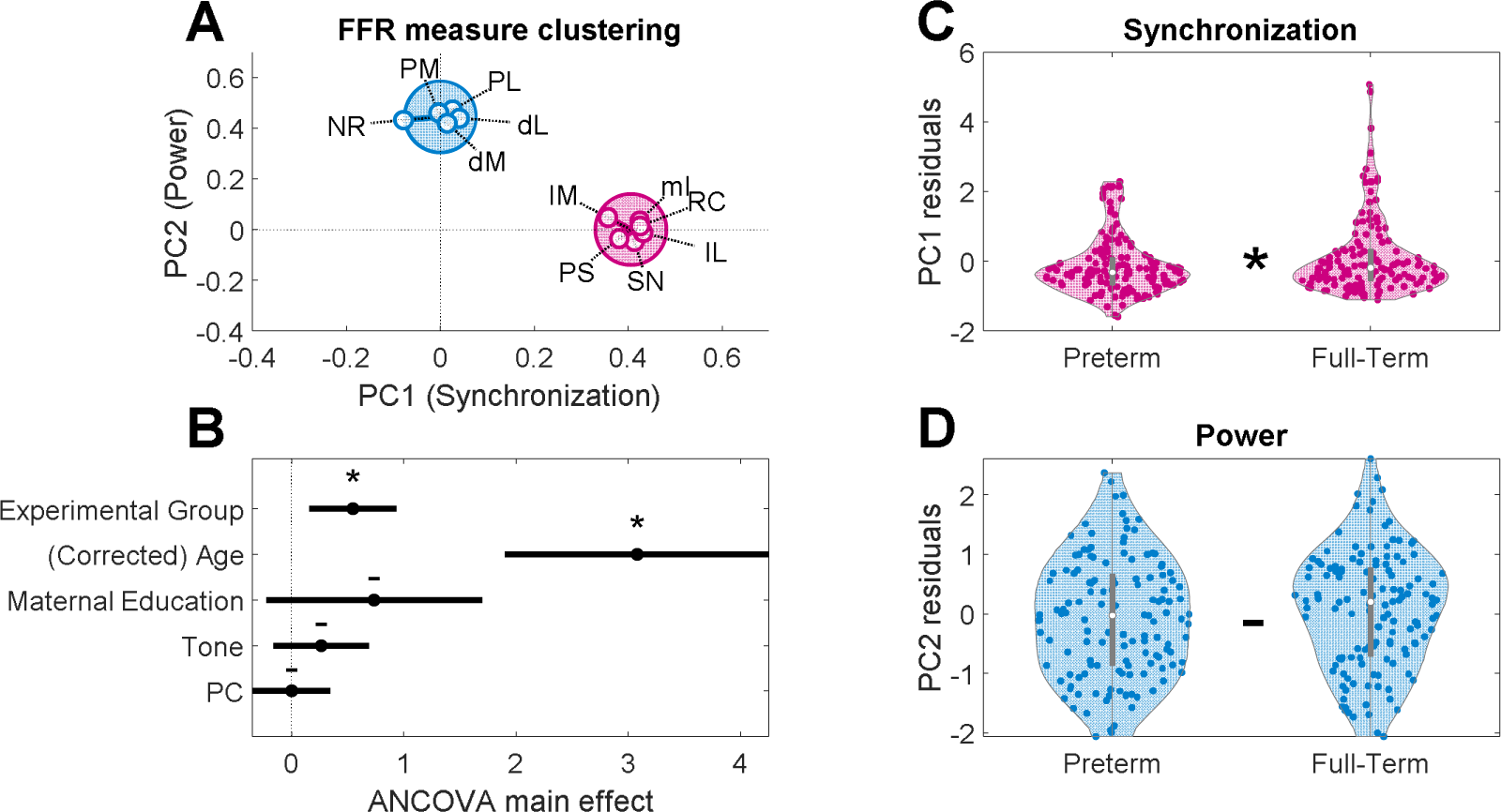
Clustering of the first two principal component coefficients of the 11 FFR measures. **(A)** The PC coefficients for each measure are plotted as empty circles connected with the lines to the centroid of their cluster that is plotted as a filled circle. Abbreviations: SN - FFR SNR, NR – Noise RMS, PL - Power LOW, PM - Power MID, mI – maxITPC, RC- Response Consistency, PS - Pitch Strength, IL - ITPC LOW, IM - ITPC MID, dL - Low/High Power Ratio, dM - Mid/High Power Ratio. AU – arbitrary units. **(B)** Main effects in ANCOVA analysis, representing an effect of one factor on the response from changing the factor value while averaging out the effects of the other factors (MathWorks, 2012). The factors are listed along the y-axes. The bars indicate the confidence intervals. The significant factors (i.e., those whose confidence intervals do not include zero) are marked with asterisks. (**C)** Synchronization and (**D**) Power findings of the two experimental groups controlled for Age, Tone, and Maternal Education. Non-parametric Wilcoxon rank-sum tests demonstrated that Synchronization (*Z* = -1.88, *P =* 0.030) but not Power (*Z* = -0.97, *P =* 0.17) was impaired by prematurity.

Dependent variables for the two groups were nonoverlapping: Synchronization dependent variables include Response Consistency (0.125±0.2011), Low-Frequency ITPC (0.053±0.0171), Mid-Frequency ITPC (0.046±0.0081), maximal ITPC (0.186±0.0748), FFR SNR (1.63±2.458 dB), and Pitch Strength (0.335±0.1296); and Power dependent variables include Low-Frequency Spectral Power (−31.5±5.161 dBmV), Mid-Frequency Spectral Power(−3138.5±3.475 dBmV), Noise RMS (0.129±0.0735 μV), Low/High Power Ratio (14.6±4.591 dB), and Mid/High Power Ratio (7.55±2.8132 dB). The numbers in parentheses indicate the mean and standard deviation of each dependent variable (note the high individual variability).

The ANCOVAs of the combined scores of the two principal components resulted in main effects of Experimental Group (*F*(1,492) = 5.11, *P* = 0.0242, η*_p_^2^ =* 0.0103) and Age (*F*(1,492) = 31.6, *P* < 0.0001, η*_p_^2^ =* 0.0603) as well as a 3-way interaction of Experimental Group x Age x Principal Component (*F*(1,492) = 14.0, *P*=0.0002, η*_p_^2^ =* 0.0277). No other main effect or interaction were significant (i.e., Tone, Maternal Education and Principal Component). The sizes of the main effects calculated by MATLAB *plotEffects* function as the effects of each factor on the response from changing the factor value while averaging out the effects of the other factors are shown at the **Figure 4B** (MathWorks, 2012).

In order to follow up the 3-way interaction, we performed two separate ANCOVAs for Synchronization and Power principal components with factors Experimental Group, Tone, Principal Component, Age, and Maternal Education. They showed that the Synchronization score was higher in full-term than in preterm infants (*F*(1,263) *=* 5.60*, P =* 0.0187, η*_p_^2^ =* 0.0208) and increased with age (*F*(1,263) = 22.62, *P =* 3.26·10^-6^, η*_p_^2^ =* 0.0792). The Power score also increased with age (*F(*1,263*) =* 11.1*, P =* 0.0010, η*_p_^2^ =* 0.0405), but by contrast did not differ between the full-term and preterm infants (*F*(1,263) *=* 0.905, *P =* 0.3424, η*_p_^2^ =* 0.0034).

Neither PC outcome differed across stimulus tones or with maternal education in years; that is, results did not depend on the nativeness of the tone or SES. Comparisons of the effects of prematurity in the two principal components are shown in **Figures 4C** and **4D**. To exclude the possibility of outliers in Synchronization data—and accordingly its deviation from normality---we performed a one-tailed non-parametric Wilcoxon rank-sum test on the sores of the two principal components controlling for age, tone, and maternal education. The test confirmed that Synchronization (*Z* = -1.88, *P =* 0.030) but not Power (*Z* = -0.97, *P =* 0.17) deteriorated with the prematurity status of the infants.

To detail which of the original Synchronization and Power FFR measures contributed most to these results, sensitivity ANCOVAs on the individual FFR measures were conducted (all *P*-values reported below are FDR-corrected, see **Table 2**). Two measures from the Synchronization cluster yielded higher values in the full-term than in the preterm infants: Response Consistency (*F*(1,264*)=*7.26*; P =* 0.0075, η*_p_^2^=* 0.0268) and maximal ITPC (*F*(1,264) *=* 10.22*; P =* 0.0016; η*_p_^2^ =* 0.0373). No Power measure showed an effect of prematurity.

All six Synchronization measures and three of five Power measures demonstrated an effect of age with EEG. The six age-dependent Synchronization measures are FFR SNR (*F*(1,264) *=* 9.40; η*_p_^2^ =* 0.0344*; P =*0.0024), Response consistency (*F(*1,264*)=* 15.04*; P =* 0.0001; η*_p_^2^ =* 0.0539), Pitch Strength (*F(*1,264*) =* 18.62*; P =* 2.25·10^-5^; η*_p_^2^ =* 0.0659), maximal ITPC (*F(*1,264*) =* 11.88*; P =* 0.0007; η *^2^ =* 0.0431), Low-Frequency ITPC (*F(*1,264*)=*18.16*; P =* 2.83·10^-5^; η*_p_^2^ =* 0.0644), and Mid-Frequency ITPC (*F(*1,264*) =* 40.75*; P =* 7.73·10^-10^; η*_p_^2^ =* 0.1337). The following Power measures improved with age: Low-Frequency Spectral Power (*F(*1,264*)=*12.40*; P =* 0.0005; η *^2^ =* 0.0449), Mid-Frequency Spectral Power (*F(*1,264*) =* 18.98*; P =* 1.90·10^-5^; η*_p_^2^ =* 0.0671), and Noise RMS (*F(*1,264*)=* 7.80*; P =* 0.0056; η*_p_^2^ =* 0.0287). Note that the effects of maturation on Low-Frequency Spectral, Mid-Frequency Spectral Power, FFR SNR, and Pitch Strength in preterm infants are consistent with earlier findings reported in term infants (Novitskiy et al., 2022).

## 4. Discussion

Advances in neonatal care have drastically increased the survival rate of preterm infants. However, preterm infants remain at risk for poorer long-term neurodevelopmental outcomes than infants born at term, including outcomes that are subtle, such as language impairment and learning disabilities. In a meta-analysis covering ∼1,500 preterm infants, Zimmerman (2018) found that at an early school age (5-9 years), children who were born before 37 weeks of gestation scored significantly worse across virtually all areas of language and literacy than their peers who were born at term. Problems in language continue to be detectable even at 12 years of age in children who were born preterm (Luu et al., 2009). More alarmingly, van Noort-van der Spek *et al*. (2012) reported that preterm- and term-born children differed most in complex language (e.g., formulating sentences) as opposed to simple language (e.g., receptive vocabulary), with differences growing significantly from 3 to 12 years of age. Even when moderate-preterm (32 < 34 weeks) and late-preterm (34 < 37 weeks) infants were studied separately from very preterm infants, poorer long-term neurodevelopmental outcomes were still observed (see McGowan et al., 2011, for a review). Importantly, language difficulties seem to be the developmental domain at greatest risk in late preterm infants. For examples, while Harris (2013) and Rabie (2015) found no increased risk for ADHD in infants who were born late preterm compared to those who were born full-term, the risks for speech and language delays were significantly elevated in those who were born late preterm. Future research should be directed to understand the biological basis of these specific developmental difficulties.

Much extant research on the nervous system of human preterm infants has focused on vascular injuries (e.g., intracranial haemorrhage; van de Bor, den Ouden, & Guit, 1992) and anatomical characteristics (Bouyssi-Kobar et al., 2016). Although empirical research can be conducted to link neuroanatomy with developmental outcomes (Ullman et al., 2015; Kelly et al., 2020), such studies do not directly link neuropathologies with cognitive functions. The present study represents an effort to identify neural functional deficits in preterm infants by investigating their neural encoding of speech. With a longitudinal study that is being conducted, we will be better positioned to link neural functional deficits with language and communication deficits.

Neural auditory responses to click (Chonchaiya et al., 2013) and speech (Wong et al., 2021) in infants are associated with early language development. However, it is not known which neural measures are disrupted in preterm infants. Here, we measured the neural encoding of speech sounds in preterm and full-term infants with the FFR, an EEG-derived non-invasive technique. We divided FFR measures into two categories: synchronization and power (**Figures 2** and **4**). We found that preterm infants experienced significant disruption of neural encoding of speech, more so with synchronization than with power, after age, SES, and other non-neural factors were controlled. Although much research (possibly including animal models) is needed to understand the neuronal processes associated with synchronization and power, we speculate that these two categories of FFR are related to myelination and synaptogenesis, respectively.

Myelination and synaptogenesis are two fundamental processes in the nervous system that peak in the perinatal period of development (Kinney & Volpe, 2017; Kostović & Jovanov-Milošević, 2006; Moore & Linthicum, 2001). Preterm birth deprives the infant brain of fatty acids and iron that are normally accrued in the last trimester of pregnancy (Georgieff & Innis, 2005; Koudelka et al., 2016; Schneider et al., 2022). This deprivation can lead to pathological changes in the activity of astrocytes and microglia that prevent pre-oligodendrocyte differentiation, which in turn results in hypomyelination (Volpe, 2019). Preterm infants also often experience perinatal hypoxia during delivery and due to cardiorespiratory pathologies (Salmaso, Jablonska, Scafidi, Vaccarino, & Gallo, 2014). Although early studies in macaques suggested that synaptogenesis develops in a programmed way and was not affected by prematurity (Bourgeois et al., 1989), in the contemporary murine model experimental hypoxia causes white matter injury (WMI) by arresting oligodendrocyte differentiation, which leads to both hypomyelination and delayed synaptogenesis (F. Wang et al., 2018). Whether and how hypomyelination and/or delayed synaptogenesis are linked to functional deficits in speech processing in preterm human infants have yet to be investigated. We speculate that FFR synchronization measures are related to myelination and FFR power measures are related to synaptogenesis. Thus, by studying these two categories of FFR measures, we take one step closer to understanding specific pathophysiological processes that underlie preterm infants’ developmental deficits.

Several factors are commonly discussed in relation to the development of FFR, including myelination, synaptic transmission, phase-locking, and tonotopicity (Anderson et al., 2015; Arenillas-Alcón et al., 2021; Jeng, Lin, & Wang, 2016), two of which, myelination and phase-locking, are intricately related. Phase-locking is achieved by synchronicity among neurons. Synchronicity over a large number of spatially dispersed neurons requires fine tuning of the conduction speed that is best achieved by myelination (Liu et al., 2006; Seidl & Rubel, 2016; Sinclair et al., 2017). Incomplete myelination can account for slower conduction and reduced synchronicity in vertebrate neural networks (Almeida & Lyons, 2017; Nave & Werner, 2021). FFR measures that indicate synchronicity are likely related to myelination and should improve with myelination in development. Phase-locking can be directly measured in EEG with inter-trial phase coherence (ITPC) which is also known as the phase-locking factor (Delorme & Makeig, 2004; Tallon-Baudry et al., 1996). This frequency-domain measure indicates phase alignment of individual trials in the epoched EEG (**Figure 2**). Maximal ITPC is used as an overall measure of phase-locking. The phase-locked response to the pitch of the voice consolidates earlier in development than the response to higher harmonics (Anderson et al., 2015; Arenillas-Alcón et al., 2021; Levi, Folsom, & Dobie, 1995) justifying separate assessment of phase-locking in these two spectral areas (ITPC LOW and ITPC MID). Another way to access phase-locking is to measure response consistency, that is the correlation in the time domain between individual trials (Skoe et al., 2015). The basic assumption of signal-averaging is that event-related neural activity, including its phase, is the same on every trial (Luck, 2005). Therefore, averaging EEG trials favors stimulus-evoked waveforms that keep their phase constant to the jittered waveforms of the same amplitude. By contrast, high-amplitude background oscillations are completely subtracted with averaging if their phases are independent of the stimulus onset. Thus, the ratio between the post-stimulus and pre-stimulus of the averaged waveform (i.e., signal-to-noise ratio or SNR) also estimates the degree of phase-locking (Anderson et al., 2015; Novitskiy et al., 2022). Pronounced peaks in autocorrelation of the averaged waveform also indicate that the signal phase is aligned between individual trials at some frequency. The maximum of autocorrelation is labelled Pitch Strength (Jeng et al., 2013, 2010). We considered these six synchronization measures (Pitch Strength, SNR, Response Consistency, maximal ITPC, Low-Frequency ITPC and Mid-Frequency ITPC) as proxies of myelination.

By contrast, measures that indicate the power of background EEG (e.g., Noise RMS) or time-locked activity (e.g., Spectral Power) are more likely to reflect the general amount of synaptic activity and should grow with increases in the number of synapses (Skoe et al., 2015). The total power of the EEG (e.g., FFR Spectral Power) reflects changes in the amount of postsynaptic activity in the brain and is likely reduced by disrupted synaptogenesis (Buzsáki, Anastassiou, & Koch, 2012; Podvalny et al., 2015). High values of pre-stimulus noise RMS in FFR in young school-age children have been interpreted to reflect an excessive number of synapses that are pruned in early adulthood (Skoe et al., 2015). Averaging EEG trials removes a substantial part of the non-phase-locked activity as described above. The remaining signal power in the pre-stimulus interval quantified as Noise RMS indicates spontaneous activity. Post-stimulus Spectral Power is a mixture of spontaneous, time-locked activity, and phase-locked activity. It can be divided into three frequency bands: low-frequencies around pitch of the voice (Power LOW), mid-frequencies in the range of the lower harmonics (Power MID), and high-frequencies for higher harmonics (Skoe et al., 2015). Besides the stimulus-related peaks, the EEG spectrum is characterised by aperiodic slope, a decrease of power from lower to the higher frequencies. This so-called 1/*f* slope is an important physiological parameter characterizing spontaneous EEG activity and synaptic dynamics behind it (Gao, Peterson, & Voytek, 2017; Podvalny et al., 2015; Voytek et al., 2015). The 1/*f* slope can be estimated by division of the Low-and Mid Frequency Power by the High-Frequency Power. We considered those five power measures (Noise RMS, Power LOW, Power MID, Low/High Power Ratio and Mid/High Power Ratio) as proxies of synaptogenesis.

A previous study demonstrated that measures of the FFR response to speech matured independently of whether the speech sound belonged to a native or non-native category of the child’s ambient linguistic environment (Novitskiy et al., 2022). In the present study, those measures were classified into two clusters based on the likelihood of their association with either myelination or synaptogenesis. Classification was first performed according to theoretical predictions about the physiological properties of the measures, followed by statistical clustering; the two approaches converged. Two measures associated with myelination, maximal ITPC and Response Consistency, were less pronounced in preterm infants (GA ≤ 34 weeks) as compared with age-and sex-matched full-term controls (with the corrected age of preterms matching the chronological age of the controls). This finding accords with results of animal studies that preterm birth is characterised by hypomyelination (Volpe, 2019). None of the five measures associated with synaptogenesis was so affected, supporting the observation in macaques that synaptogenesis develops in a programmed way and is not affected by prematurity (Bourgeois et al., 1989). As found previously (Novitskiy et al., 2022), none of these findings could be attributed to whether the speech sound encoded by the infant was native or non-native.

Earlier studies demonstrated a negative relation between the SES of families and the prevalence of preterm birth, with maternal education being the most common SES measure (see McHale et al., 2022, and Wong & Edwards, 2013, for systematic reviews). Maternal education is also associated with preterm birth in three large-scale cohort studies (Poulsen et al., 2019). However, effects of maternal education on prematurity vary between countries (Ruiz et al., 2015). In the present study maternal education did not differ between preterm and full-term groups and no effect of maternal education was found in our statistical models. The plausible explanation may be overall high maternal education in our sample (median of 16 years) and relative homogeneity of the groups in terms of SES. Nor did SES contribute to predicting native language outcome above EEG responses in a previous study in Hong Kong (Wong et al., 2021).

Although the study-level results reported here are consistent with the interpretation of hypomyelination in preterm infants relative to full-term infants, it is important to underscore the substantial variability that was evident at the individual child level. In a previous study of mostly full-term infants, individual differences in FFR measures predicted future child language development (Wong et al., 2021). Individual differences may be meaningful as well in preterm infants (Putnick et al., 2017). Longitudinal language outcome data are being collected from these preterm and term children to construct a predictive algorithm for future language development that relies on individual differences in the FFR measures. Neural encoding of speech may well serve as an important augmentation of the current click ABR test (Richard et al., 2020) used to screen for hearing and language impairment and delay. Early screening will enable clinicians to prescribe preventive early intervention to potentially mitigate the extent and burden of widespread language impairment in children born preterm.

## 5. Limitations

Although the broader goal of the present study is to explore links between neural and language development in preterm children, longitudinal language outcome data are not yet available in these children. This circumstance substantially limits some conclusions of our study. Because the relation between neural encoding of speech and language outcome was established in a previous study of full-term infants (Wong et al., 2021), we have no reason to believe that such a relation would not exist in preterm infants as well. Nevertheless, preterm infants may develop compensatory mechanisms in other neural pathways to overcome their reduced neural encoding precision in order to develop language in the typical range. Future studies should evaluate the relation between neural encoding of speech and language outcomes in preterm infants empirically. Another limitation of the present study is that no direct measures of myelination were obtained. The relation between neural timing and white matter measures has only been established in cortical long-latency studies, as far as we are aware (see Babaeeghazvini, Rueda-Delgado, Gooijers, Swinnen, & Daffertshofer, 2021, for a review). Future research should connect FFR synchronicity measures with white matter neuroimaging metrics to more directly examine their association. As a study comparing preterm and full-term infants, our focus was on group-level differences. Our sample size did not allow us to investigate gestational age as a continuous variable. A future study with a large sample size should investigate more specifically the impact of gestational age on neural encoding of speech.

## 6. Conclusions

Early intervention to remediate symptoms of language impairment in children at-risk for communication disorders is effective (Roberts, Curtis, Sone, & Hampton, 2019). Preterm birth can lead to subtle language and cognitive deficits that may not surface until the preschool years. Not every preterm infant will develop these deficits. A prognostic tool is needed to differentiate those preterm infants who may or may not develop these problems so that intervention can be implemented at the earliest time point specifically for children who are at-risk for these problems. Results from the present study may lead to such a prognostic tool in the future. Importantly, because our results pointed to a subset of FFR features to be especially impaired, more weight may be put to these features to increase the tool’s precision.

## Abbreviations

ABR: auditory brainstem response
ANCOVA: analysis of covariance
FA: fractional anisotropy
FDR: False-discovery rate
FFR: frequency-following response
ISI: inter-stimulus interval
ITPC: inter-trial phase coherence
LLRs: long-latency responses
RMS: root-mean square
SES: socioeconomic status
SNR: signal-to-noise ratio
SOA: stimulus-onset-asynchrony
TF: time-frequency
WMI: white matter injury

## Figure legends

## Tables

**Table 1.**
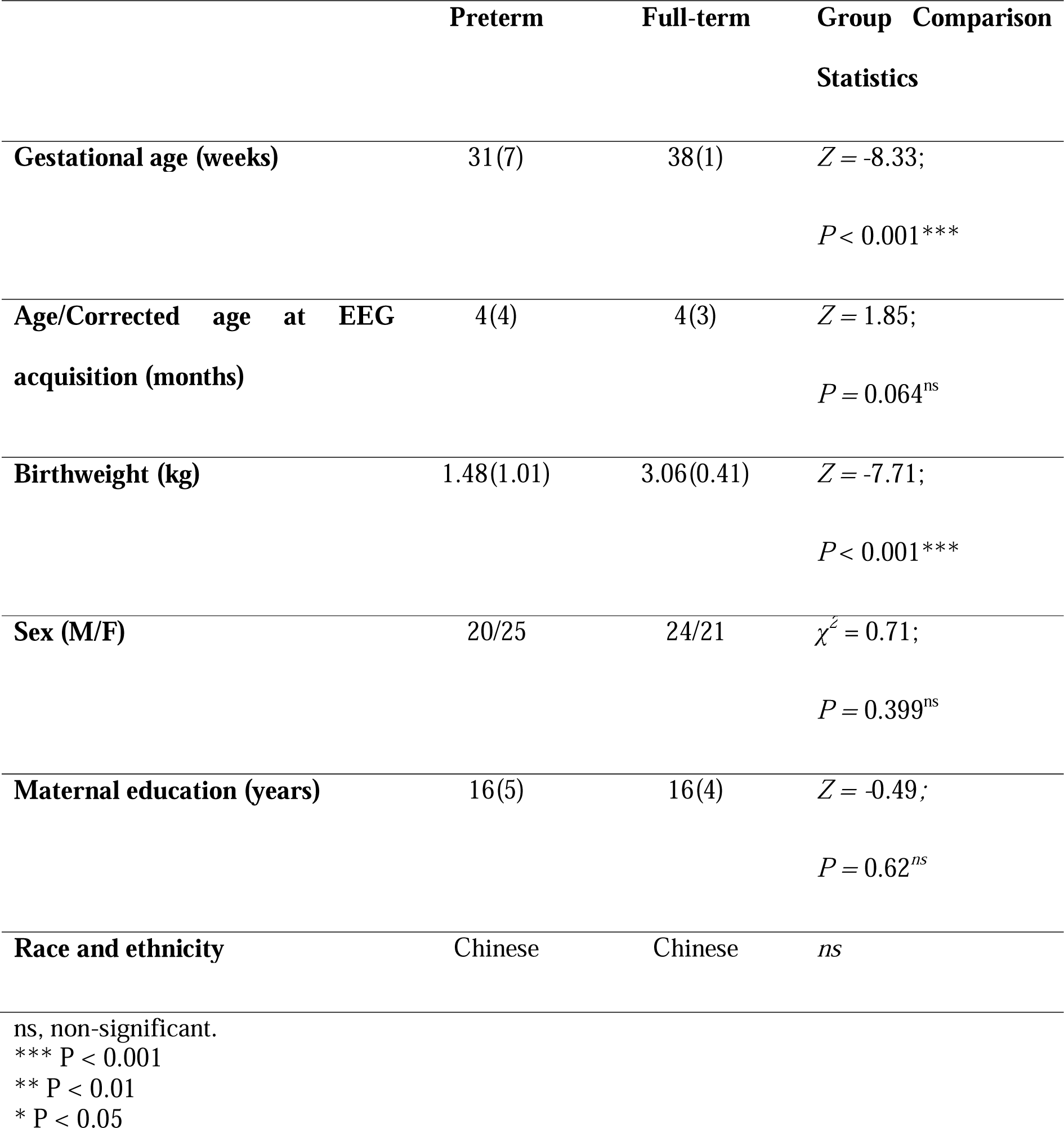
Basic sociodemographic characteristics of the infant participants (median and interquartile range)

|  | Preterm | Full-term | Group Comparison Statistics |
| --- | --- | --- | --- |
| **Gestational age (weeks)** | 31(7) | 38(1) | $Z = -8.33$;<br>$P < 0.001^{***}$ |
| **Age/Corrected age at EEG acquisition (months)** | 4(4) | 4(3) | $Z = 1.85$;<br>$P = 0.064^{ns}$ |
| **Birthweight (kg)** | 1.48(1.01) | 3.06(0.41) | $Z = -7.71$;<br>$P < 0.001^{***}$ |
| **Sex (M/F)** | 20/25 | 24/21 | $\chi^2 = 0.71$;<br>$P = 0.399^{ns}$ |
| **Maternal education (years)** | 16(5) | 16(4) | $Z = -0.49$;<br>$P = 0.62^{ns}$ |
| **Race and ethnicity** | Chinese | Chinese | $ns$ |

ns, non-significant.
\*\*\* $P < 0.001$
\*\* $P < 0.01$
\* $P < 0.05$

**Table 2.**
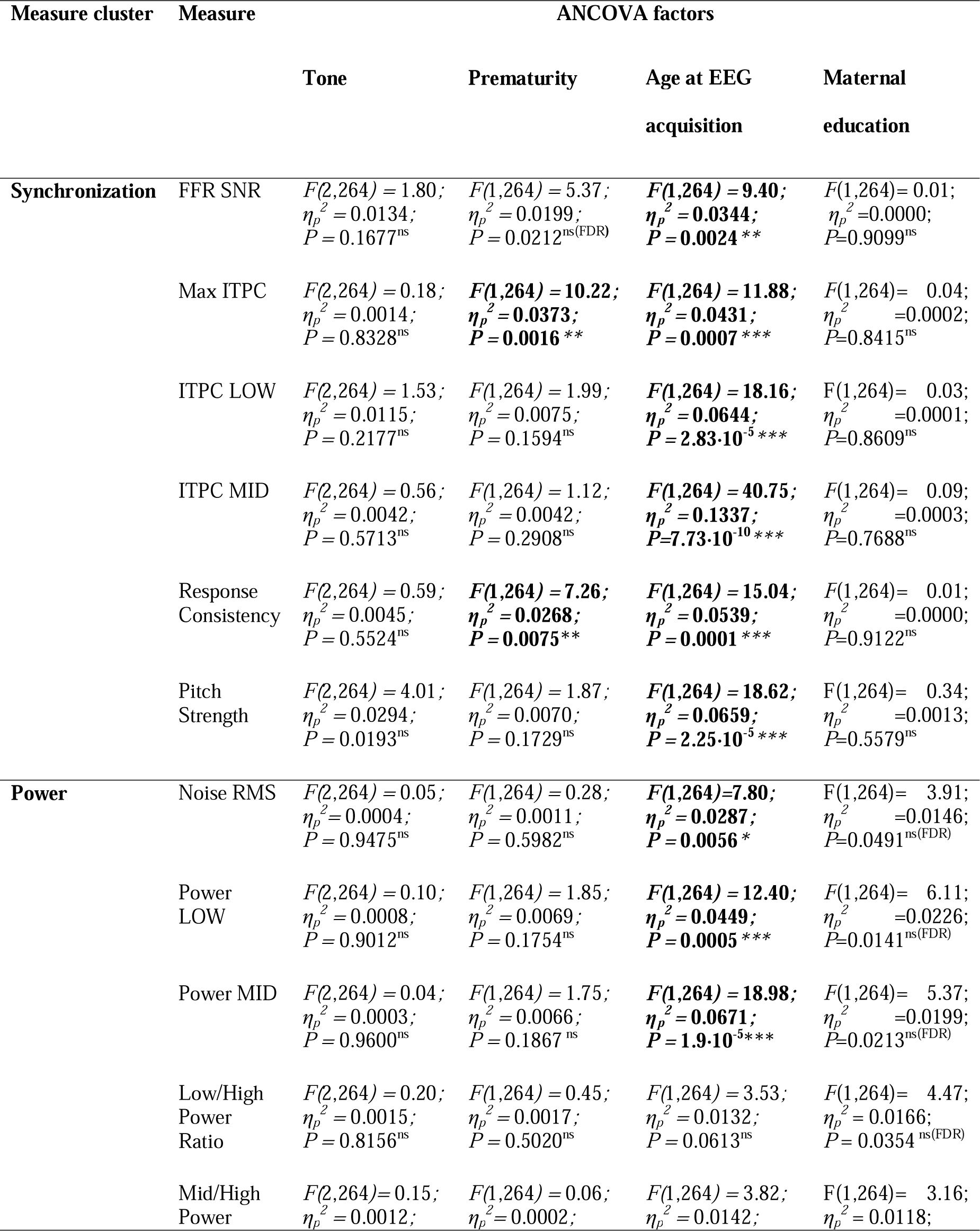
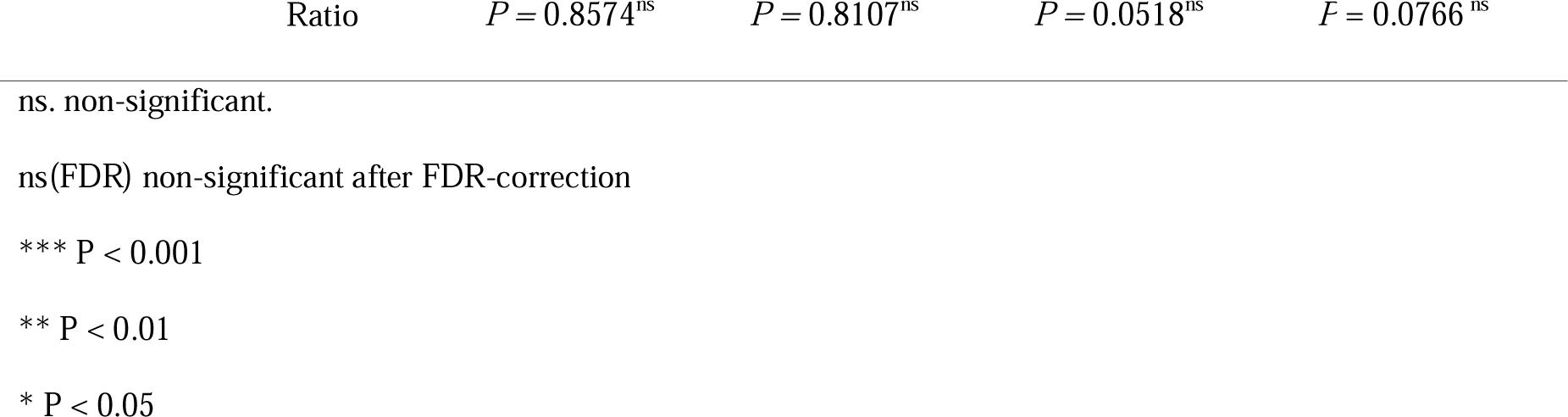
Results of the 3-way fixed-effects ANCOVA for individual measures (FDR-corrected across measures and factors)

## References

1. Almeida, R. G., & Lyons, D. A. (2017). On myelinated axon plasticity and neuronal circuit formation and function. The Journal of Neuroscience, 37(42), 10023–10034. https://doi.org/10.1523/JNEUROSCI.3185-16.2017

2. Anderson, S., Parbery-Clark, A., White-Schwoch, T., & Kraus, N. (2012). Aging affects neural precision of speech encoding. Journal of Neuroscience, 32(41), 14156–14164. https://doi.org/10.1523/JNEUROSCI.2176-12.2012

3. Anderson, S., Parbery-Clark, A., White-Schwoch, T., & Kraus, N. (2015). Development of subcortical speech representation in human infants. The Journal of the Acoustical Society of America, 137(6), 3346–3355. https://doi.org/10.1121/1.4921032

4. Arenillas-Alcón, S., Costa-Faidella, J., Ribas-Prats, T., Gómez-Roig, M. D., & Escera, C. (2021). Neural encoding of voice pitch and formant structure at birth as revealed by frequency-following responses. Scientific Reports, 11, 6660. https://doi.org/10.1038/s41598-021-85799-x

5. Babaeeghazvini, P., Rueda-Delgado, L. M., Gooijers, J., Swinnen, S. P., & Daffertshofer, A. (2021). Brain Structural and Functional Connectivity: A Review of Combined Works of Diffusion Magnetic Resonance Imaging and Electro-Encephalography. Frontiers in Human Neuroscience, 15, 585. https://doi.org/10.3389/FNHUM.2021.721206/BIBTEX

6. Benjamini, Y., & Hochberg, Y. (1995). Controlling the false discovery rate: a practical and powerful approach to multiple testing. Journal of the Royal Statistical Society. Series B (Methodological).

7. Blencowe, H., Cousens, S., Oestergaard, M. Z., Chou, D., Moller, A.-B., Narwal, R., … Lawn, J. E. (2012). National, regional, and worldwide estimates of preterm birth rates in the year 2010 with time trends since 1990 for selected countries: a systematic analysis and implications. The Lancet, 379(9832), 2162–2172. https://doi.org/10.1016/S0140-6736(12)60820-4

8. Bornstein, M. H. (2019). A developmentalist’s viewpoint: “It’s about time!” Ecological systems, transaction, and specificity as key developmental principles in children’s changing worlds. In Children in Changing Worlds (pp. 277–286). Cambridge University Press. https://doi.org/10.1017/9781108264846.010

9. Bornstein, M. H., Hahn, C.-S., Suwalsky, J. T. D., & Haynes, O. M. (2003). Socioeconomic Status, Parenting, and Child Development: The Hollingshead Four-Factor Index of Social Status and the Socioeconomic Index of Occupations. In M. H. Bornstein & R. H. Bradley (Eds.), Socioeconomic status, parenting, and child development (pp. 29–82). Routledge. https://doi.org/10.4324/9781410607027

10. Bourgeois, J.-P., Jastreboff, P. J., & Rakic, P. (1989). Synaptogenesis in visual cortex of normal and preterm monkeys: evidence for intrinsic regulation of synaptic overproduction. Proceedings of the National Academy of Sciences, 86(11), 4297–4301. https://doi.org/10.1073/pnas.86.11.4297

11. Bouyssi-Kobar, M., du Plessis, A. J., McCarter, R., Brossard-Racine, M., Murnick, J., Tinkleman, L., … Limperopoulos, C. (2016). Third trimester brain growth in preterm infants compared with in utero healthy fetuses. Pediatrics, 138(5), e20161640. https://doi.org/10.1097/01.ogx.0000513225.92648.a4

12. Brignoni-Pérez, E., Dubner, S. E., Ben-Shachar, M., Berman, S., Mezer, A. A., Feldman, H. M., & Travis, K. E. (2022). White matter properties underlying reading abilities differ in 8-year-old children born full term and preterm: A multi-modal approach. NeuroImage, 256. https://doi.org/10.1016/J.NEUROIMAGE.2022.119240

13. Buzsáki, G., Anastassiou, C. A., & Koch, C. (2012). The origin of extracellular fields and currents — EEG, ECoG, LFP and spikes. Nature Reviews Neuroscience, 13(6), 407–420. https://doi.org/10.1038/nrn3241

14. Chonchaiya, W., Tardif, T., Mai, X., Xu, L., Li, M., Kaciroti, N., … Lozoff, B. (2013). Developmental trends in auditory processing can provide early predictions of language acquisition in young infants. Developmental Science, 16(2), 159–172. https://doi.org/10.1111/desc.12012

15. Cohen, M. X. (2017). MATLAB for Brain and Cognitive Scientists. The MIT Press.

16. Committee on fetus and newborn. (2004). Age terminology during the perinatal period. PEDIATRICS, 114(5), 1362–1364. https://doi.org/10.1542/peds.2004-1915

17. Delorme, A., & Makeig, S. (2004). EEGLAB: An open source toolbox for analysis of single-trial EEG dynamics including independent component analysis. Journal of Neuroscience Methods, 134(1), 9–21. https://doi.org/10.1016/j.jneumeth.2003.10.009

18. Dibble, M., Ang, J. Z., Mariga, L., Molloy, E. J., & Bokde, A. L. W. (2021). Diffusion tensor imaging in very preterm, moderate-late preterm and term-born neonates: a systematic review. The Journal of Pediatrics, 232, 48–58.e3. https://doi.org/10.1016/j.jpeds.2021.01.008

19. Eimas, P. D., Siqueland, E. R., Jusczyk, P., & Vigorito, J. (1971). Speech perception in infants. *Science (New York*, N.Y.), 171, 303–306.

20. Font-Alaminos, M., Cornella, M., Costa-Faidella, J., Hervás, A., Leung, S., Rueda, I., & Escera, C. (2020). Increased subcortical neural responses to repeating auditory stimulation in children with autism spectrum disorder. Biological Psychology, 149, 107807. https://doi.org/10.1016/j.biopsycho.2019.107807

21. Gao, R., Peterson, E. J., & Voytek, B. (2017). Inferring synaptic excitation/inhibition balance from field potentials. NeuroImage, 158, 70–78. https://doi.org/10.1016/j.neuroimage.2017.06.078

22. Georgieff, M. K., & Innis, S. M. (2005). Controversial nutrients that potentially affect preterm neurodevelopment: essential fatty acids and iron. Pediatric Research, 57, 99R-103R. https://doi.org/10.1203/01.PDR.0000160542.69840.0F

23. Guit, G. L., van de Bor, M., den Ouden, L., & Wondergem, J. H. M. (1990). Prediction of neurodevelopmental outcome in the preterm infant: MR-staged myelination compared with cranial US. Radiology, 175(1), 107–109. https://doi.org/10.1148/radiology.175.1.2179986

24. Harris, M. N., Voigt, R. G., Barbaresi, W. J., Voge, G. A., Killian, J. M., Weaver, A. L., … Katusic, S. K. (2013). ADHD and Learning Disabilities in Former Late Preterm Infants: A Population-Based Birth Cohort. Pediatrics, 132(3), e630–e636. https://doi.org/10.1542/peds.2012-3588

25. Hoormann, J., Falkenstein, M., Hohnsbein, J., & Blanke, L. (1992). The human frequency-following response (FFR): Normal variability and relation to the click-evoked brainstem response. Hearing Research, 59(2), 179–188. https://doi.org/10.1016/0378-5955(92)90114-3

26. Jeng, F.-C., Lin, C.-D., Sabol, J. T., Hollister, G. R., Chou, M.-S., Chen, C.-H., … Tsou, Y.-A. (2016). Pitch perception and frequency-following responses elicited by lexical-tone chimeras. International Journal of Audiology, 55(1), 53–63. https://doi.org/10.3109/14992027.2015.1072774

27. Jeng, F.-C., Lin, C.-D., & Wang, T.-C. (2016). Subcortical neural representation to Mandarin pitch contours in American and Chinese newborns. The Journal of the Acoustical Society of America, 139(6), EL190–EL195. https://doi.org/10.1121/1.4953998

28. Jeng, F.-C., Lin, C.-D. Der, Chou, M. S., Hollister, G. R., Sabol, J. T., Mayhugh, G. N., … Wang, C.-Y. Y. (2016). Development of subcortical pitch representation in three-month-old Chinese infants. Perceptual and Motor Skills, 122(1), 123–135. https://doi.org/10.1177/0031512516631054

29. Jeng, F.-C., Peris, K. S., Hu, J., & Lin, C.-D. (2013). Evaluation of an automated procedure for detecting frequency-following responses in American and Chinese neonates. Perceptual and Motor Skills, 116(2), 456–465. https://doi.org/10.2466/24.10.PMS.116.2.456-465

30. Jeng, F.-C., Schnabel, E. A., Dickman, B. M., Hu, J., Li, X., Lin, C.-D., & Chung, H.-K. (2010). Early maturation of frequency-following responses to voice pitch in infants with normal hearing. Perceptual and Motor Skills, 111(3), 765–784. https://doi.org/10.2466/10.22.24.PMS.111.6.765-784

31. Kelly, C. E., Thompson, D. K., Cheong, J. L. Y., Chen, J., Olsen, J. E., Eeles, A. L., … Spittle, A. J. (2019). Brain structure and neurological and behavioural functioning in infants born preterm. Developmental Medicine & Child Neurology, 61(7), 820–831. https://doi.org/10.1111/dmcn.14084

32. Kidokoro, H., Anderson, P. J., Doyle, L. W., Woodward, L. J., Neil, J. J., & Inder, T. E. (2014). Brain injury and altered brain growth in preterm infants: predictors and prognosis. Pediatrics, 134(2), e444–e453. https://doi.org/10.1542/peds.2013-2336

33. Kinney, H. C., & Volpe, J. J. (2017). Myelination events. In Volpe’s Neurology of the Newborn (Sixth Edit, Vol. 13, pp. 176–188). Elsevier Inc. https://doi.org/10.1016/B978-0-323-42876-7.00008-9

34. Kostović, I., & Jovanov-Milošević, N. (2006). The development of cerebral connections during the first 20-45 weeks’ gestation. Seminars in Fetal and Neonatal Medicine, 11(6), 415–422. https://doi.org/10.1016/j.siny.2006.07.001

35. Koudelka, S., Voas, M. G. M., Almeida, R. G., Baraban, M., Soetaert, J., Meyer, M. M. P., … Lyons, D. A. (2016). Individual neuronal subtypes exhibit diversity in CNS myelination mediated by synaptic vesicle release. Curr Biol, 26(11), 1447–1455. https://doi.org/10.1016/j.cub.2016.03.070

36. Kraus, N., & White-Schwoch, T. (2020). Listening in on the listening brain. The Hearing Journal, 73(7), 46. https://doi.org/10.1097/01.HJ.0000689460.50136.1d

37. Krizman, J., & Kraus, N. (2019). Analyzing the FFR: A tutorial for decoding the richness of auditory function. Hearing Research, 382, 107779. https://doi.org/10.1016/j.heares.2019.107779

38. Lerner, R. M., & Bornstein, M. H. (2021). Contributions of the specificity principle to theory, research, and application in the study of human development: A view of the issues. Journal of Applied Developmental Psychology, 75, 101294. https://doi.org/10.1016/j.appdev.2021.101294

39. Levi, E. C., Folsom, R. C., & Dobie, R. A. (1995). Coherence analysis of envelope-following responses (EFRs) and frequency-following responses (FFRs) in infants and adults. Hearing Research, 89(1–2), 21–27. https://doi.org/10.1016/0378-5955(95)00118-3

40. Liu, L.-F., Palmer, A. R., & Wallace, M. N. (2006). Phase-locked responses to pure tones in the inferior colliculus. Journal of Neurophysiology, 95(3), 1926–1935. https://doi.org/10.1152/jn.00497.2005

41. Luck, S. J. (2005). An Introduction to the Event-related Potential Technique. MIT Press. https://doi.org/10.4155/fmc.12.40

42. Luu, T. M., Ment, L. R., Schneider, K. C., Katz, K. H., Allan, W. C., & Vohr, B. R. (2009). Lasting Effects of Preterm Birth and Neonatal Brain Hemorrhage at 12 Years of Age. Pediatrics, 123(3), 1037–1044. https://doi.org/10.1542/peds.2008-1162

43. MathWorks. (2012). plotEffects. Retrieved from https://ww2.mathworks.cn/help/stats/linearmodel.ploteffects.html

44. McGowan, J. E., Alderdice, F. A., Holmes, V. A., & Johnston, L. (2011). Early Childhood Development of Late-Preterm Infants: A Systematic Review. Pediatrics, 127(6), 1111– 1124. https://doi.org/10.1542/peds.2010-2257

45. McHale, P., Maudsley, G., Pennington, A., Schlüter, D. K., Barr, B., Paranjothy, S., & Taylor-Robinson, D. (2022). Mediators of socioeconomic inequalities in preterm birth: a systematic review. BMC Public Health, 22(1), 1–20. https://doi.org/10.1186/s12889-022-13438-9

46. Moore, J. K., & Linthicum, F. H. (2001). Myelination of the human auditory nerve: different time courses for schwann cell and glial myelin. *Annals of Otology*, Rhinology & Laryngology, 110(7), 655–661. https://doi.org/10.1177/000348940111000711

47. Munakata, S., Okada, T., Okahashi, A., Yoshikawa, K., Usukura, Y., Makimoto, M., … Okuhata, Y. (2013). Gray matter volumetric MRI differences late-preterm and term infants. Brain and Development, 35(1), 10–16. https://doi.org/10.1016/j.braindev.2011.12.011

48. Nave, K.-A., & Werner, H. B. (2021). Ensheathment and myelination of axons: evolution of glial functions. Annual Review of Neuroscience, 44(1), 197–219. https://doi.org/10.1146/annurev-neuro-100120-122621

49. Novitskiy, N., Maggu, A. R., Lai, C. M., Chan, P. H. Y., Wong, K. H. Y., Lam, H. S., … Wong, P. C. M. (2022). Early development of neural speech encoding depends on age but not native language status: evidence from lexical tone. Neurobiology of Language, 3(1), 67–86. https://doi.org/10.1162/nol_a_00049

50. Podvalny, E., Noy, N., Harel, M., Bickel, S., Chechik, G., Schroeder, C. E., … Malach, R. (2015). A unifying principle underlying the extracellular field potential spectral responses in the human cortex. Journal of Neurophysiology, 114(1), 505–519. https://doi.org/10.1152/jn.00943.2014

51. Polka, L., & Werker, J. F. (1994). Developmental changes in perception of nonnative vowel contrasts. Journal of Experimental Psychology: Human Perception and Performance, 20(2), 421–435. https://doi.org/10.1037/0096-1523.20.2.421

52. Poulsen, G., Andersen, A.-M. N., Jaddoe, V. W. V, Magnus, P., Raat, H., Stoltenberg, C., … Mortensen, L. H. (2019). Does smoking during pregnancy mediate educational disparities in preterm delivery? Findings from three large birth cohorts. Paediatric and Perinatal Epidemiology, 33(2), 164–171. https://doi.org/10.1111/ppe.12544

53. Putnick, D. L., Bornstein, M. H., Eryigit-Madzwamuse, S., & Wolke, D. (2017). Long-term stability of language performance in very preterm, moderate-late preterm, and term children. The Journal of Pediatrics, 181, 74–79.e3. https://doi.org/10.1016/j.jpeds.2016.09.006

54. Rabie, N. Z., Bird, T. M., Magann, E. F., Hall, R. W., & McKelvey, S. S. (2015). ADHD and developmental speech/language disorders in late preterm, early term and term infants. Journal of Perinatology, 35(8), 660–664. https://doi.org/10.1038/jp.2015.28

55. Ribas[Prats, T., Arenillas-Alcón, S., Lip[Sosa, D. L., Costa[Faidella, J., Mazarico, E., Gómez[Roig, M. D., & Escera, C. (2022). Deficient neural encoding of speech sounds in term neonates born after fetal growth restriction. Developmental Science, 25, e13189. https://doi.org/10.1111/desc.13189

56. Ribeiro, F. M., & Carvallo, R. M. (2008). Tone-evoked ABR in full-term and preterm neonates with normal hearing. International Journal of Audiology, 47(1), 21–29. https://doi.org/10.1080/14992020701643800

57. Richard, C., Neel, M. L., Jeanvoine, A., Connell, S. M., Gehred, A., Maitre, N. L., … Maitre, N. L. (2020). Characteristics of the Frequency-Following Response to Speech in Neonates and Potential Applicability in Clinical Practice: A Systematic Review. Journal of Speech, Language, and Hearing Research, 63(5), 1618–1635. https://doi.org/10.1044/2020_JSLHR-19-00322

58. Roberts, M. Y., Curtis, P. R., Sone, B. J., & Hampton, L. H. (2019). Association of Parent Training with Child Language Development: A Systematic Review and Meta-analysis. JAMA Pediatrics. https://doi.org/10.1001/jamapediatrics.2019.1197

59. Ruiz, M., Goldblatt, P., Morrison, J., Kukla, L., Švancara, J., Riitta-Järvelin, M., … Pikhart, H. (2015). Mother’s education and the risk of preterm and small for gestational age birth: a DRIVERS meta-analysis of 12 European cohorts. Journal of Epidemiology and Community Health, 69(9), 826–833. https://doi.org/10.1136/jech-2014-205387

60. Salmaso, N., Jablonska, B., Scafidi, J., Vaccarino, F. M., & Gallo, V. (2014). Neurobiology of premature brain injury. Nature Neuroscience, 17(3), 341–346. https://doi.org/10.1038/nn.3604

61. Sansavini, A., Guarini, A., Justice, L. M., Savini, S., Broccoli, S., Alessandroni, R., & Faldella, G. (2010). Does preterm birth increase a child’s risk for language impairment? Early Human Development, 86(12), 765–772. https://doi.org/10.1016/j.earlhumdev.2010.08.014

62. Schlegel, A. A., Rudelson, J. J., & Tse, P. U. (2012). White matter structure changes as adults learn a second language. Journal of Cognitive Neuroscience, 24(8), 1664–1670. https://doi.org/10.1162/jocn_a_00240

63. Schneider, N., Bruchhage, M. M. K., O’Neill, B. V., Hartweg, M., Tanguy, J., Steiner, P., … Deoni, S. C. L. (2022). A nutrient formulation affects developmental myelination in term infants: a randomized clinical trial. Frontiers in Nutrition, 9, 823893. https://doi.org/10.3389/fnut.2022.823893

64. Seidl, A. H., & Rubel, E. W. (2016). Systematic and differential myelination of axon collaterals in the mammalian auditory brainstem. Glia, 64(4), 487–494. https://doi.org/10.1002/glia.22941

65. Silva, D., Lopez, P., & Mantovani, J. (2014). Auditory brainstem response in term and preterm infants with neonatal complications: the importance of the sequential evaluation. International Archives of Otorhinolaryngology, 19, 161–165. https://doi.org/10.1055/s-0034-1378137

66. Sinclair, J. L., Fischl, M. J., Alexandrova, O., Heβ, M., Grothe, B., Leibold, C., & Kopp-Scheinpflug, C. (2017). Sound-evoked activity influences myelination of brainstem axons in the trapezoid body. The Journal of Neuroscience, 37(34), 8239–8255. https://doi.org/10.1523/JNEUROSCI.3728-16.2017

67. Skoe, E., & Kraus, N. (2010). Auditory brainstem reponse to complex sounds[: a tutorial. Ear and HearingHear, 31(3), 302–324. https://doi.org/10.1097/AUD.0b013e3181cdb272.Auditory

68. Skoe, E., Krizman, J., Anderson, S., & Kraus, N. (2015). Stability and plasticity of auditory brainstem function across the lifespan. Cerebral Cortex, 25(6), 1415–1426. https://doi.org/10.1093/cercor/bht311

69. Sleifer, P., da Costa, S. S., Cóser, P. L., Goldani, M. Z., Dornelles, C., & Weiss, K. (2007). Auditory brainstem response in premature and full-term children. International Journal of Pediatric Otorhinolaryngology, 71(9), 1449–1456. https://doi.org/10.1016/j.ijporl.2007.05.029

70. Stipdonk, L. W., Weisglas-Kuperus, N., Franken, M.-C. J., Nasserinejad, K., Dudink, J., & Goedegebure, A. (2016). Auditory brainstem maturation in normal-hearing infants born preterm: a meta-analysis. Developmental Medicine & Child Neurology, 58(10), 1009–1015. https://doi.org/10.1111/dmcn.13151

71. Tallon-Baudry, C., Bertrand, O., Delpuech, C., & Pernier, J. (1996). Stimulus specificity of phase-locked and non-phase-locked 40 Hz visual responses in human. The Journal of Neuroscience, 16(13), 4240–4249. https://doi.org/10.1523/JNEUROSCI.16-13-04240.1996

72. The American college of obstetricians and gynecologists committee. (2013). Committee opinion 579: definition of term pregnancy. OBSTETRICS & GYNECOLOGY, 122(5), 248–251. https://doi.org/10.1097/SPV.0000000000000113

73. Thompson, D. K., Kelly, C. E., Chen, J., Beare, R., Alexander, B., Seal, M. L., … Spittle, A. J. (2019). Characterisation of brain volume and microstructure at term-equivalent age in infants born across the gestational age spectrum. NeuroImage: Clinical, 21, 101630. https://doi.org/10.1016/j.nicl.2018.101630

74. Ullman, H., Spencer-Smith, M., Thompson, D. K., Doyle, L. W., Inder, T. E., Anderson, P. J., & Klingberg, T. (2015). Neonatal MRI is associated with future cognition and academic achievement in preterm children. Brain, 138(11), 3251–3262. https://doi.org/10.1093/brain/awv244

75. van de Bor, M., den Ouden, L., & Guit, G. L. (1992). Value of cranial ultrasound and magnetic resonance imaging in predicting neurodevelopmental outcome in preterm infants. Pediatrics, 90(2), 196–199. https://doi.org/10.1542/peds.90.2.196

76. van Noort-van der Spek, I. L., Franken, M.-C. J. P., & Weisglas-Kuperus, N. (2012). Language Functions in Preterm-Born Children: A Systematic Review and Meta-analysis. Pediatrics, 129(4), 745–754. https://doi.org/10.1542/peds.2011-1728

77. Volpe, J. J. (2019). Dysmaturation of premature brain: importance, cellular mechanisms, and potential interventions. Pediatric Neurology, 95, 42–66. https://doi.org/10.1016/j.pediatrneurol.2019.02.016

78. Voytek, B., Kramer, M. A., Case, J., Lepage, K. Q., Tempesta, Z. R., Knight, R. T., & Gazzaley, A. (2015). Age-related changes in 1/f neural electrophysiological noise. Journal of Neuroscience, 35(38), 13257–13265. https://doi.org/10.1523/JNEUROSCI.2332-14.2015

79. Walani, S. R. (2020). Global burden of preterm birth. International Journal of Gynecology & Obstetrics, 150(1), 31–33. https://doi.org/10.1002/ijgo.13195

80. Walsh, J. M., Doyle, L. W., Anderson, P. J., Lee, K. J., & Cheong, J. L. Y. (2014). Moderate and late preterm birth: effect on brain size and maturation at term-equivalent age. Radiology, 273(1), 232–240. https://doi.org/10.1148/radiol.14132410

81. Wang, F., Yang, Y.-J., Yang, N., Chen, X.-J., Huang, N.-X., Zhang, J., … Mei, F. (2018). Enhancing oligodendrocyte myelination rescues synaptic loss and improves functional recovery after chronic hypoxia. Neuron, 99, 689–701. https://doi.org/10.1016/j.neuron.2018.07.017

82. Wang, W., Yu, Q., Liang, W., Xu, F., Li, Z., Tang, Y., & Liu, S. (2022). Altered cortical microstructure in preterm infants at term-equivalent age relative to term-born neonates. Cerebral Cortex, in press. https://doi.org/10.1093/cercor/bhac091

83. Wong, H. S., & Edwards, P. (2013). Nature or Nurture: A Systematic Review of the Effect of Socio-economic Status on the Developmental and Cognitive Outcomes of Children Born Preterm. Maternal and Child Health Journal, 17(9), 1689–1700. https://doi.org/10.1007/s10995-012-1183-8

84. Wong, P. C. M., Lai, C. M., Chan, P. H. Y., Leung, T. F., Lam, H. S., Feng, G., … Novitskiy, N. (2021). Neural speech encoding in infancy predicts future language and communication difficulties. American Journal of Speech-Language Pathology, 30(5), 2241–2250. https://doi.org/10.1044/2021_AJSLP-21-00077

85. Woodward, L. J., Moor, S., Hood, K. M., Champion, P. R., Foster-Cohen, S., Inder, T. E., & Austin, N. C. (2009). Very preterm children show impairments across multiple neurodevelopmental domains by age 4 years. Archives of Disease in Childhood: Fetal and Neonatal Edition, 94(5), F339–44. https://doi.org/10.1136/adc.2008.146282

86. Zimmerman, E. (2018). Do Infants Born Very Premature and Who Have Very Low Birth Weight Catch Up With Their Full Term Peers in Their Language Abilities by Early School Age? *Journal of Speech*, Language, and Hearing Research, 61(1), 53–65. https://doi.org/10.1044/2017_JSLHR-L-16-0150

